# *In vitro* and *in vivo* efficacy of the combination of colistin and endolysins against clinical strains of Multi-Drug Resistant (MDR) pathogens

**DOI:** 10.1101/662460

**Authors:** L. Blasco, A. Ambroa, R. Trastoy, E. Perez-Nadales, F. Fernández-Cuenca, J. Torre-Cisneros, J. Oteo, A. Oliver, R. Canton, T. Kidd, F. Navarro, E. Miró, A. Pascual, G. Bou, L. Martínez-Martínez, M. Tomas, GEMARA SEIMC/REIPI Bacterial Clinical Adaptation Study Group

## Abstract

The multidrug resistance (MDR) among pathogenic bacteria is jeopardizing the worth of antimicrobials, which had previously changed medical sciences. In this study, we used bioinformatic tools to identify the endolysins ElyA1 and ElyA2 (GH108-PG3 family) present in the genome of bacteriophages Ab1051Φ and Ab1052Φ, respectively. The muralytic activity of these endolysins over MDR clinical isolates (*Acinetobacter baumannii*, *Pseudomonas aeruginosa* and *Klebsiella pneumoniae*) was tested using the turbidity reduction assay. The minimal inhibitory concentrations (MICs) of endolysin, colistin and their combination were determined using the microdilution checkerboard method. The antimicrobial activity of the combinations was confirmed by time kill curves and *in vivo* assays in larvae of *Galleria mellonella*. Our results showed that ElyA1 displayed activity against all 25 strains of *A. baumannii* and *P. aeruginosa* tested and against 13 out of 17 strains of *K. pneumoniae*. No activity was detected when assays were done with endolysin ElyA2. The combined antimicrobial activity of colistin and endolysin ElyA1 yielded a reduction in the colistin MIC for all strains studied, except *K. pneumoniae*. These results were confirmed *in vivo* in *G. mellonella* survival assays. In conclusion, the combination of colistin with new endolysins such as ElyA1 could increase the bactericidal activity and reduce the MIC of the antibiotic, thus also reducing the associated toxicity.

**IMPORTANCE:** The development of multiresistance by pathogen bacteria increases the necessity of the development of new antimicrobial strategies. In this work, we combined the effect of the colistin with a new endolysin, ElyA1, from a bacteriophage present in the clinical strain of *Acinetobacter baumannii* Ab105. ElyA1 is a lysozyme-like family (GH108-GP3), whose antimicrobial activity was described for first time in this work. Also, another endolysin, ElyA2, with the same origin and family, was characterized but in this case no activity was detected. ElyA1 presented lytic activity over a broad spectrum of strains from *A. baumannii, Pseudomonas aeruginosa*, and *Klebsiella pneumoniae*. When colistin was combined with ElyA1 an increase of the antimicrobial activity was observed with a reduced concentration of colistin, and this observation was also confirmed *in vivo* in *Galleria mellonella* larvae. The combination of colistin with new endolysins as ElyA1 could increase the bactericidal activity and lowering the MIC of the antibiotic, thus also reducing the associated toxicity.

## INTRODUCTION

The worldwide emergence of multidrug resistant (MDR) microorganisms that are refractory to treatment with current therapeutic agents has emphasised the urgent need for new classes of antimicrobial agents (1). Recently, the World Health Organization (WHO) published a list of “priority pathogens” which includes those microorganisms that are considered a serious threat to human health and for which new anti-infective treatments are urgently needed. The members of this list are carbapenem-resistant *A.baumannii*, *P.aeruginosa* and *K.pneumoniae* clinical isolates (2).

A consequence of the emergence of the MDR bacteria is the return to the use of antimicrobials whose use was reduced or abandoned. This was the case of colistin or polymixin E, a cationic peptide which disturbs the stability of the outer membrane increasing its permeability through electrostatic interactions and cationic displacement of the lipopolysaccharide. In spite of its antimicrobial effects, this antibiotic presented nephrotoxicity effects and finally its use was gradually dismissed and substituted by other more tolerable antibiotics (3, 4). The search of new antimicrobial agents as well as their combination with old antibiotics such as colistin are new strategies in seeking novel treatments against MDR microorganisms.

In recent years, a novel drug discovery approach has explored endolysin enzymes (also referred to as enzybiotics) encoded by bacteriophages (viruses which infect bacteria) (5). Endolysins are actively produced during the lytic cycle and exert antibacterial activity through degradation of peptidoglycan in the bacterial cell wall (5, 6).

Endolysins are highly evolved enzymes produced by bacteriophages to digest the bacterial cell wall at the end of their replication cycle and release the phage progeny. Endolysins target the integrity of the cell wall and attack one of the major bonds in the peptidoglycan layer. They can be classified into five groups according to the cleavage site: N-acetyl-β-D-muramidase (lysozymes); N-acetyl-β-D-glucosaminidases (glycosidases); lytic transglycosylase; N-acetylmuramoyl-L-alanine amidases and L-alanoyl-D-glutamate endopeptidases (6, 7).

Endolysins are good candidates as new antimicrobial agents against Gram-positive bacteria, in which the peptidoglycan layer of the cell wall is exposed to the medium. Several studies have evaluated the potential use of endolysin against Gram-positive bacteria such as *Staphylococcus aureus, Streptococcus agalactiae*, *Streptococcus pneumoniae* and *Streptococcus pyogenes* in animal models of human infections and diseases (8–16). In Gram-negative bacteria, the outer membrane acts as a barrier to many endolysins, and very few endolysins with exogenous activity against Gram-negative bacteria have been described (many are biotechnologically engineered) (17–20).

Endolysins can attack Gram-negative bacteria when the outer membrane is previously permeabilized with agents such as EDTA, which destabilizes the lipopolysaccharides of the outer membrane; however, the combination of endolysin and EDTA is limited to a topical treatment of localized infections (21, 22). In the search for alternative methods of killing MDR bacteria such as *A. baumannii*, *P.aeruginosa* and *K.pneumoniae* various researchers have considered increasing the muralytic activity of endolysins by combining them with different antibiotics to take advantage of synergistic responses (22–25).

In this report, we identified and characterized an endolysin, named ElyA1, isolated from the *A. baumannii* Ab105 (ROC0034a) bacteriophage Ab1051Φ. The endolysin displayed muralytic activity against a broad spectrum of MDR organisms. In addition, combining endolysin ElyA1 with colistin (polymyxin E) enhanced the susceptibility of the tested strains to colistin by at least four times, thus highlighting the potential of endolysin ElyA1 as an antibacterial agent candidate. This effect was confirmed by a test *in vivo*, in which the survival of the *G. mellonella* larvae increased when colistin (¼ MIC) was supplemented with endolysin ElyA1. Moreover, another endolysin from the same family, named ElyA2, was identified in the *A. baumannii* Ab105 bacteriophage Ab1052Φ, but no muralytic activity was detected in this enzyme.

## MATERIALS AND METHODS

### Strains and culture conditions

The bacterial strains and plasmids used in this study included 25 *A. baumannii* MDR strains belonging to 22 different sequence types (STs) (Table 1). The strains were isolated from colonized or infected patients within the framework of the II Spanish Multicentre Study which counted with the participation of 45 Spanish hospitals (GEIH-REIPI *Acinetobacter baumannii* 2000-2010, Genbank Umbrella Bioproject accession number PRJNA422585)(26). They included 25 MDR clinical strains of *P. aeruginosa* (many included in CC274), all of which were isolated from cystic fibrosis patients, and 17 carbapenemase-producing strains of *K. pneumoniae*, which were isolated in 20 Spanish hospitals during the EuSCAPE project (27, 28). Moreover, *Escherichia coli* DH5α and Rosetta strains were used in cloning assays (Table 1).

**Table 1.**
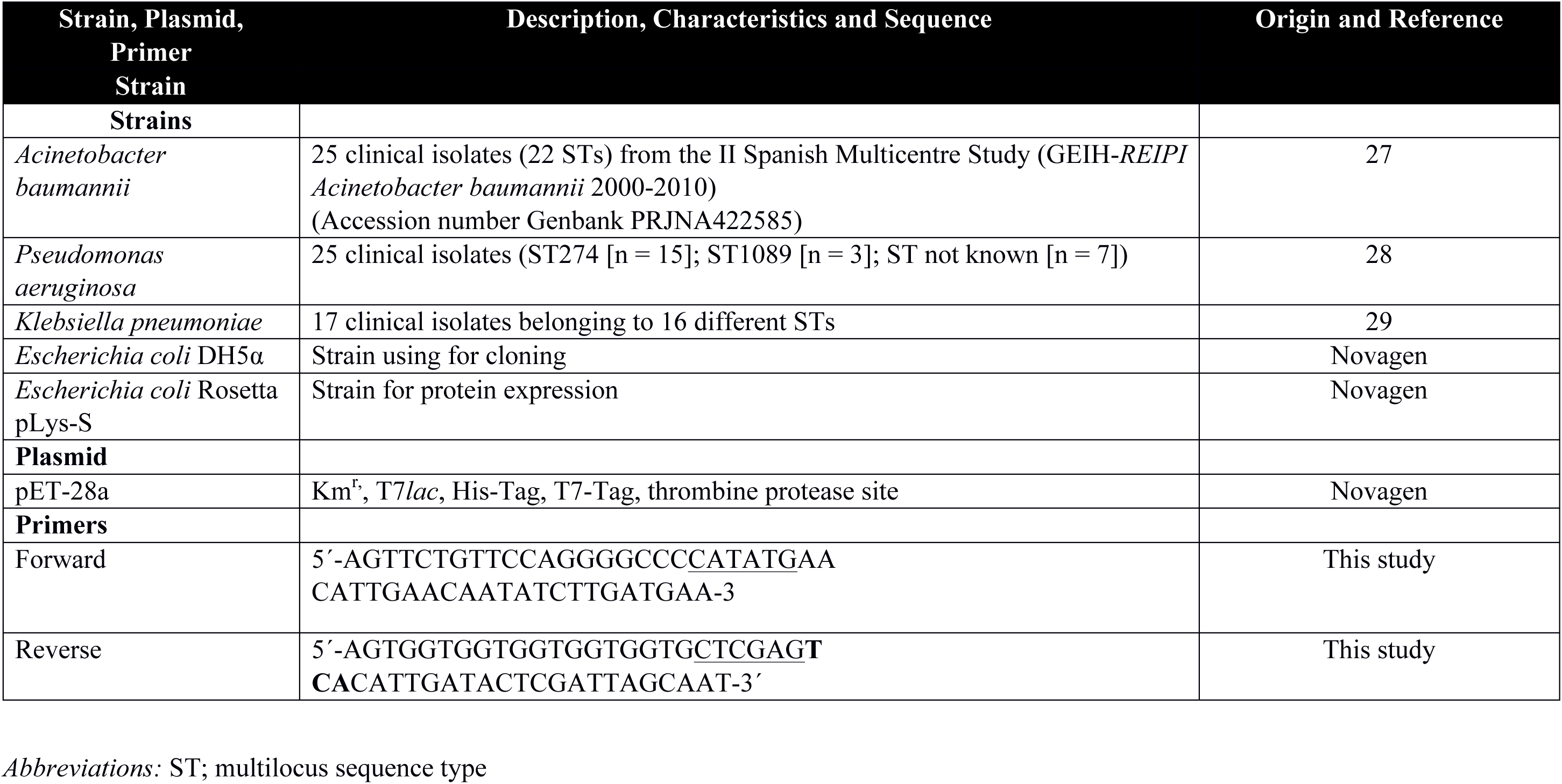
Description of the bacterial strains, plasmids and primers used in this study.

All strains were cultured in LB (Luria Bertani) at 180 rpm and 37°C. For solid medium, 2% of agar was added to LB broth. When transformation assays were done the medium was supplemented with 50μg/ml of ampicillin.

### Identification and Purification of the endolysins ElyA1 and ElyA2

Endolysin gene prediction, from the genome of the bacteriophage Ab105Φ1 (GenBank: KT588074.1) and Ab105Φ2 (GenBank: KT588075.2) (26) (Figure 1), was performed with the bioinformatic tools PHASTER (Phage Search Tool Enhanced Release) and RAST (Rapid Annotation Using Subsystem). Protein homology analysis was performed by BLAST (Basic Local Alignment Search Tool), Clustal Omega and MView. Protein families were assigned using InterProScan and the domain graphic was assigned with PROSITE MyDomains.

**Figure 1.**
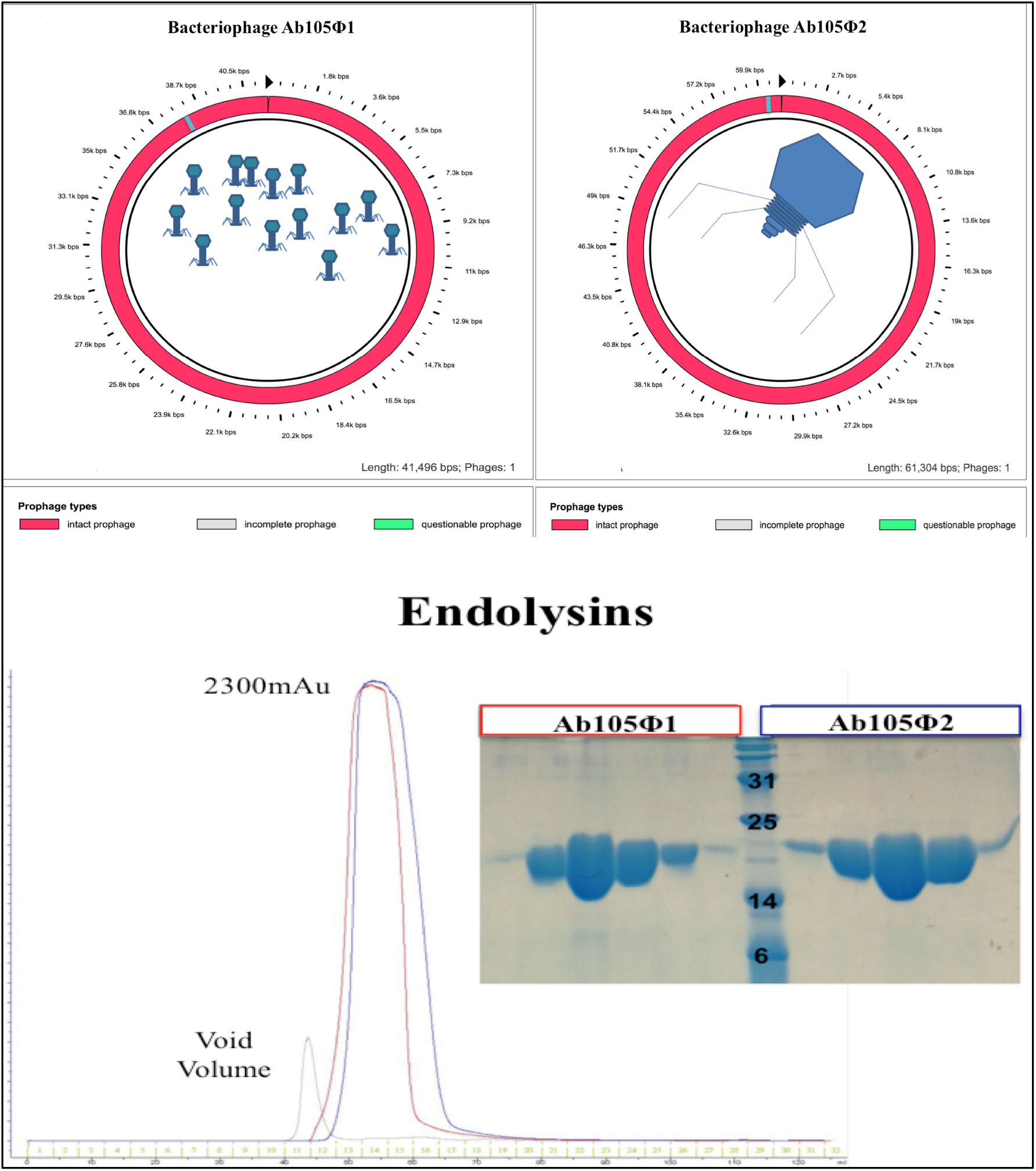
Genome of the bacteriophages Ab105Φ1 (GenBank: KT588074.1) and Ab105Φ2 (GenBank: KT588075.2) by figure modified from PHAST software.. SDS-PAGE purification of the endolysins ElyA1 and ElyA2 (chromatography study).

The endolysin genes were amplified by PCR from the genomic DNA of *A. baumannii* Ab105, which contains the DNA of the prophages Ab105Φ1 and Ab105Φ2, and cloned into the expression vector pET-28a (Novagen). The recombinant plasmids were transformed into competent *E. coli* DH5α cells (Novagen) for DNA production and purification, and the integrity of both constructs was verified by sequencing. All of the primers used are listed in Table 1. Finally, the plasmids were transformed into *Escherichia coli* Rosetta pLys-S cells (Novagen) for expression of the protein.

After induction with 1 mM IPTG, the culture (1 l) was grown at 30°C for 5 h. The bacterial cells were recovered by centrifugation (in a JLA 81000 rotor, Beckman-Coulter, at 6 Krpm for 15 min) and disrupted by sonication (VibraCell 75042 sonicator, Bioblock Scientific, tip model CV33). The sample was centrifuged in a JA 25-50 rotor (Beckman-Coulter) at 20 Krpm for 30 min. The recovered supernatant was filtered using 0.45 μm syringe-driven filters (Jet Biofil) and loaded in a His-Trap column (GE Healthcare) equilibrated with 350 mM NaCl, 50 mM Tris pH 7.5, 1 mM TCEP and 10 mM Imidazole. The proteins were eluted with 350 mM NaCl, 50 mM Tris pH 7.5, 1 mM TCEP and 150 mM Imidazole. After concentration in an Amicon Ultracel 10,000 MCWO concentrator (Millipore), the sample was loaded into a Superdex 75 16/60 column (GE Healthcare), equilibrated with 150 mM NaCl, 20 mM Tris pH 7.5 and 1 mM TCEP. The protein was eluted in a single peak. Finally, the pooled peak fractions were concentrated to 40 mg/ml, as previously described. The purification process was carried out at 4° C, and the purity was determined by SDS-PAGE (Figure 1).

### Determination of the muralytic activity of endolysins ElyA1 and ElyA2

Muralytic activity was determined using the Gram-negative overlay method described by Schmitz et al (29). Briefly, two clinical isolates of *A. baumannii*, MAR001 and PAU002, were grown to stationary phase (10^9^ CFU/ml) in LB, pelleted and resuspended in PBS buffer pH 7.4. Agar was added directly to the bacteria suspension suspension at a concentration of 0.8% and autoclaved for 15min at 120°C. The medium, containing the disorganized cells with the peptidoglycan exposed, was solidified in Petri dishes and aliquots (50 μg) of endolysin, or the endolysin buffer as negative control were spotted on the surface.

The muralytic activity was measured using as target a culture of *A. baumannii* Ab105 treated with EDTA in order to permeabilize the outer membrane. An overnight culture of *A. baumannii* Ab105 was diluted 1:100 in LB medium and grown to exponential phase (0.3-0.4 OD600nm). The culture was centrifuged (3000g, 10min), and the resulting pellet was resuspended in 20mM Tris-HCl buffer at pH 8.5 with 0.5 mM EDTA before being incubated for 30 min at room temperature. The pellet was recovered by centrifugation and washed twice in Tris-HCl buffer pH 8.5. Finally, the cells were resuspended in 20 mM Tris-HCl 150 mM NaCl pH8.5 and 25 μg/ml of endolysin ElyA1. The activity was measured by the turbidity reduction assay, as a decrease in the OD600nm after incubation with after with shaking at 37°C (17). The optical density was measured at intervals of 5 minutes for a period of 20 minutes and the time point of the higher activity was established. Also, the optimal pH and temperature for the endolysin activity were determined employing the turbidity reduction assay. The reaction was done as described previously, with the Tris-HCl at different pH (range 6.5 to 9) and temperature (room temperature, 30°C and 37°C).

### Antibacterial assays

The antibacterial activity of the endolysin was assayed with all of the 67 clinical strains of *A. baumannii, P. aeruginosa* and *K. pneumoniae* (Table 1). The activity was determined using the turbidity reduction assay, as previously described, at pH 8.5 and 37°C. The incubation times in the presence of EDTA varied according to the species assayed: 30 min for *A. baumannii* and *K. pneumoniae* and 15 min for *P. pneumoniae.*

### Broth microdilution checkerboard assay and microdilution test to determine minimum inhibitory concentrations (MICs)

This assay was done in those strains with the higher and the lower susceptibility to endolysin. All the strains tested were susceptible to colistin except three strains that were colistin resistant: *A. baumannii* SOF004b, *P. aeruginosa* AUS034 and *K. pneumoniae* KP2. The effect of the interaction between endolysin and colistin was done by the chequerboard MIC assay. In a 96-well microtiter plate with Mueller-Hinton Broth (MHB), 7 serially double dilutions were done for endolysin and 6 for colistin. The wells were then inoculated with the test culture at a final concentration of 10^5^ colony forming units (cfu/ml). The MIC of colistin (0 to 2 μgml^−1^) and concentration of endolysin ElyA1 protein (3.125 to 200 μgml^−1^) was assayed independently in the same plate. The MIC was determined as the concentration of antimicrobial agent in the well where no visible growth of bacteria was observed after incubation for 24h at 35°C. The MIC_AB_ was defined as the MIC of drug A in the presence of drug B, and the MIC_BA_ was defined as the MIC of drug B in the presence of drug A.

### Time kill curve assay

Time kill curve assays were carried out with those strains in which the MIC of colistin in ElyA1-colistin combinations was decreased by at least fourfold in checkerboard assays. The assay was conducted according to previously described techniques (30). Flasks of LB containing colistin and colistin plus endolysin at the concentration corresponding to the checkerboard assay were inoculated with a dilution 1:100 of an overnight culture in stationary phase of the tested strain and incubated at 37°C and 180 rpm in a shaking incubator. Aliquots were removed after 0, 6 and 24 h and were serially diluted and plated to produce colony forming units (cfu). Synergy was established when a ≤2 −log_10_ decrease in cells counts at 6 or 24 h in the antimicrobial combination relative to the most active single agent was observed. No effect was considered to have occurred when the counts were <2−log_10_ lower or higher relative to the culture with the single agent. Antagonism was defined when the counts in the culture with antimicrobial combination were ≥ 2−log_10_ higher than in the culture with single most active antimicrobial agent.

The reduction in the MIC of the colistin in combination with endolysins was also assayed by combining colistin with another endolysin ElyA2, isolated from the bacteriophage Ab105Φ2. The curve was done for the same strains and under the same conditions as for colistin-ElyA1.

### *Galleria mellonella* Infection Model

The *Galleria mellonella* model was an adapted version of that developed by Peleg et al (31, 32). The procedure was as follows: twelve *G. mellonella* larvae, acquired from TruLarvTM (Biosystems Technology, Exeter, Devon, UK), were each injected with 10 μl of a suspension of *A. baumannii* PON001, diluted in sterile phosphate buffer saline (PBS) containing 1 × 10^5^ CFU (± 0. 5 log). The injection was performed with a Hamilton syringe (volume 100 μl) (Hamilton, Shanghai, China). An hour after infection, larvae were injected with 10 μl of colistin in combination with endolysin ElyA1, colistin in combination with endolysin ELyA2, and colistin alone as a control group, all at the concentrations used in the time kill curve. After being injected, the larvae were placed in Petri dishes and incubated in darkness at 37°C. The number of dead larvae was recorded for 5 days. The larvae were considered dead when they showed no movement in response to touch (31).

The mortality curves corresponding to the *in vivo Galleria mellonella* infection model were constructed using GraphPad Prism v.6 and the data were analysed using the Graham-Breslow-Wilcoxon test. In both cases, p-values <0.05 were considered statistically significant, and the data were expressed as mean values.

## RESULTS

### Identification of endolysins ElyA1 and ElyA2

The gene of 546 bp coding for endolysin ElyA1 was identified as an ORF (Open Reading Frame) encoding a protein of 181 aa (GenBank: ALJ99090.1) and 20.22 kDa. The protein sequence was analysed with InterProScan and classified as a lysozyme (N-acetylmuramidase) with a C-terminal domain corresponding with the Glycosyde hydrolase 108 superfamily and also a Peptidoglycan Binding domain PG3 at the N-terminal end.

Protein homology analysis revealed a high level of homology (more than 80%) with a group of 9 endolysins from *A. baumannii* bacteriophages belonging to the same protein family as ElyA1 (20). Endolysin ElyA2 gene of 543 bp was identified as an ORF encoding a protein of 180 aa (GenBank: ALJ99174.1) and with a MW of 20.19 kDa. The sequence analysis revealed that ElyA2 protein is also a lysozyme (N-acetylmuramidase) with a C-terminal domain corresponding with the Glycosyde hydrolase 108 superfamily and also a Peptidoglycan Binding domain PG3 at the N-terminal end.

As ElyA1 protein, this enzyme presented a high homology (more than 80%) with the same group of 9 endolysins and also a homology of 90% with ElyA1 protein (20).

### Endolysins muralytic activity characterization

In the first screening of the muralytic activity of the purified endolysin ElyA1 in the Gram-negative overlay plates, a lysis halo appeared in both strains of *A. baumannii* tested (Figure 2a).

**Figure 2.**
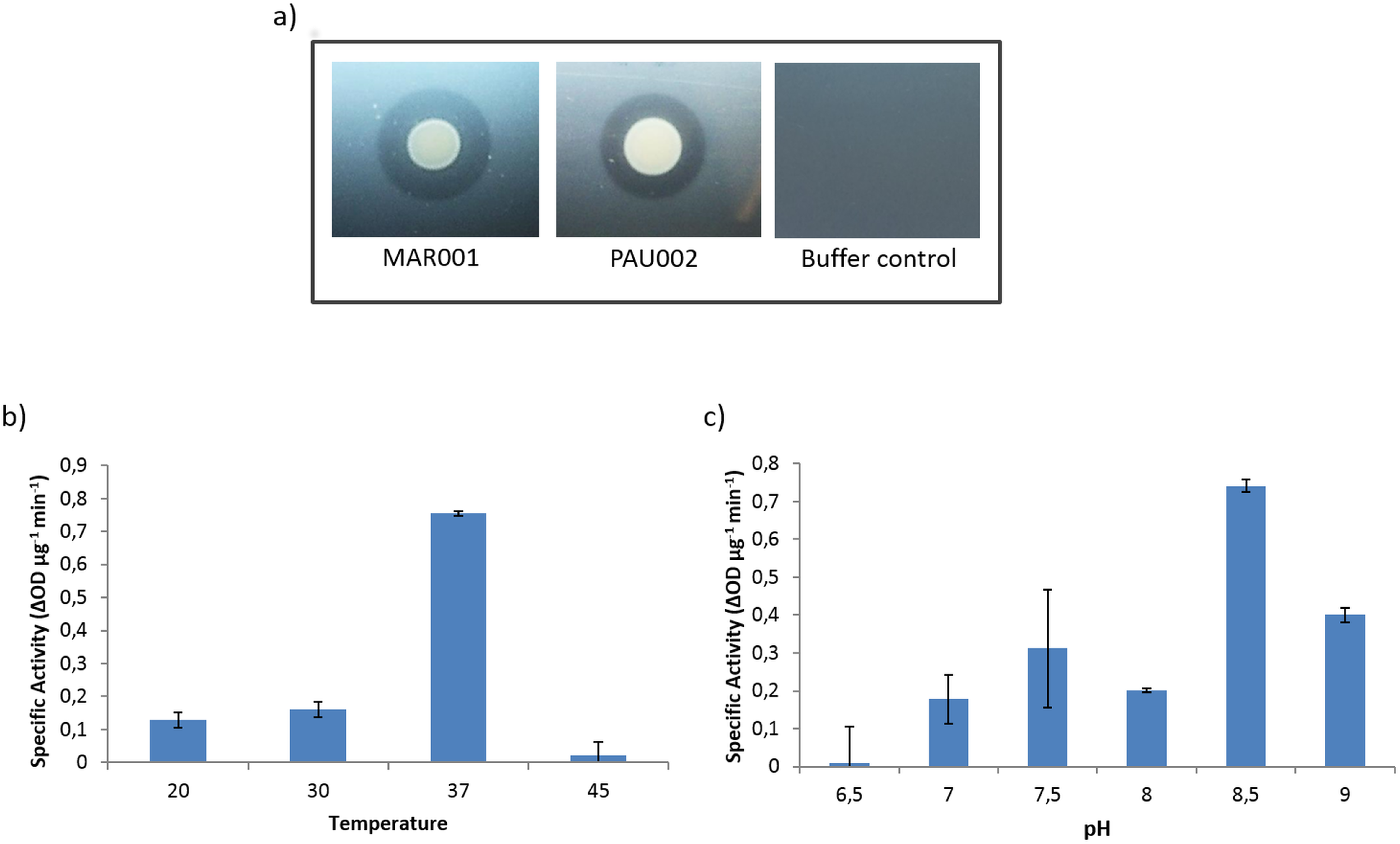
Characterization of enzymatic activity: a) Muralytic activity of ElyA1 was done by spots of ElyA1 and endolysin buffer as negative control in a Gram-negative overlay of two *Acinetobacter baumannii* clinical isolates, MAR001 and PAU002; b) pH range and c) Temperature range, were determined by the specific activity measured as the difference in optical density of the culture per μg of enzyme and minute.

The muralytic activity of this enzyme was characterized using the Gram-negative bacteria *A. baumannii* Ab105 as substrate, as this is the host strain for the phage Ab105Φ1. The enzymatic activity was measured after different incubation at different temperatures and pH. The maximum activity was obtained after incubation for 10 min at pH8.5 and 37°C (Figure 2b-c). In addition, the muralytic activity on the cells of Ab105 was assayed directly or after treatment of the cells with EDTA to permeabilize the outer membrane. However, no activity was detected when the enzyme was added directly to the cells whose outer membrane was not previously permeabilized with EDTA, and also in this case the cells tended to aggregate (data not shown).

The antibacterial assays showed a broad lytic spectrum of activity against the strains of the three species tested (Figure 3). As expected because of the origin of the *A. baumannii* endolysin, the highest activity observed was among the 25 *A. baumannii* strains tested. Although the activity was more variable in *P. aeruginosa*, muralytic activity was detected against all of the strains tested. Finally, endolysin ElyA1 displayed activity against 13 of the 17 *K. pneumoniae* strains, although at lower levels than in *A. baumannii* and *P. aeruginosa*. The strains tested in the three species belonged to different STs, but no correlation between the susceptibility to endolysin ElyA1 and the ST was detected.

**Figure 3.**
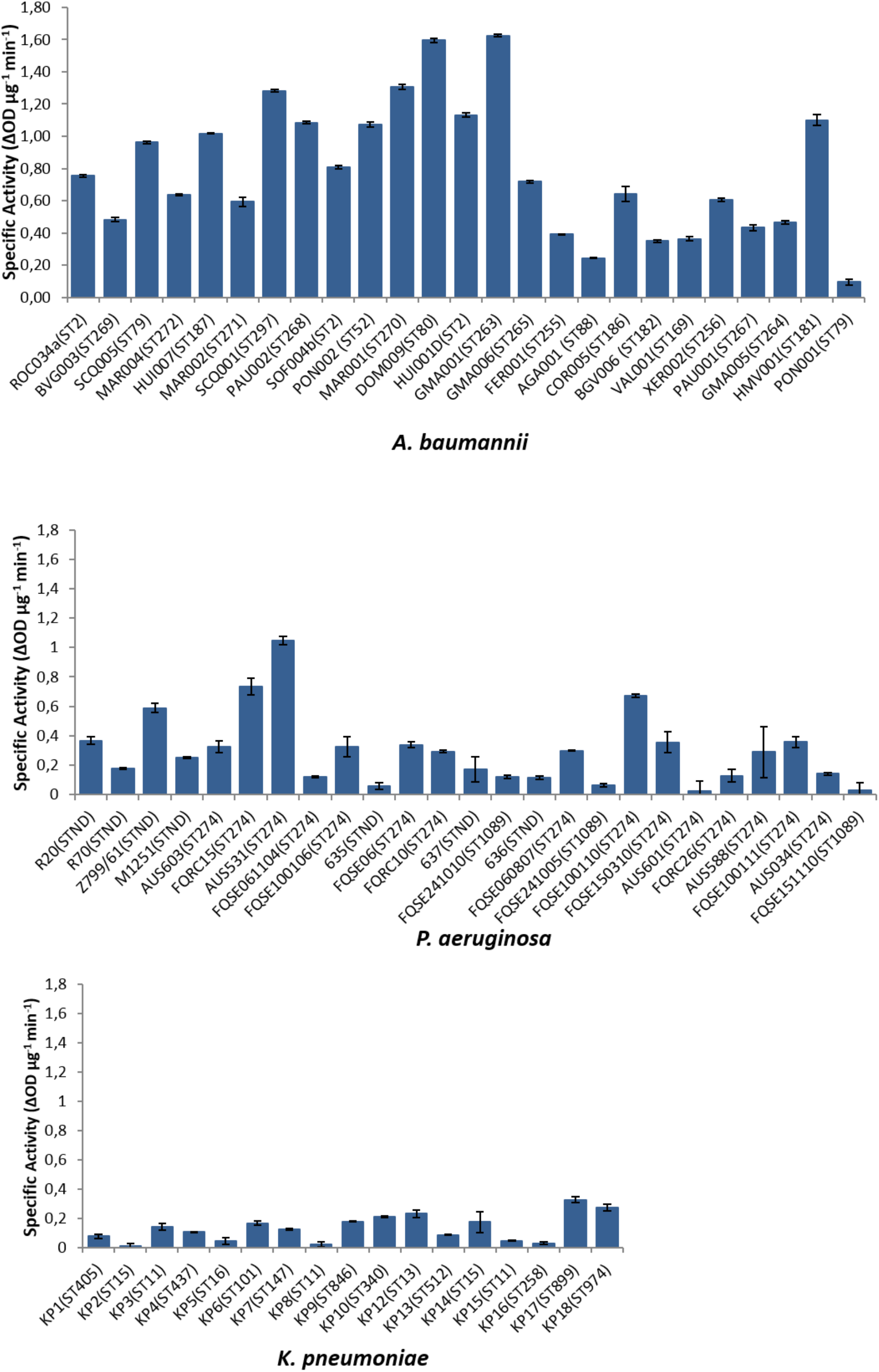
Specific activity of endolysin ElyA1 tested in clinical isolates from different mutlilocus sequence types (STs) of three Gram-negative members of the ESKAPE group: *Acinetobacter. baumannii, Pseudomonas aeruginosa and Klebsiella pneumoniae*.

When muralytic activity of endolysin ElyA2 was tested no activity was detected under any condition assayed. On the contrary, this enzyme induces the aggregation of the cells at all the enzyme concentration tested both in those cells treated previously with EDTA and in those with an intact outer membrane.

### Endolysin ElyA1 and colistin combination activity Assays

As endolysin ElyA1 is only active when the outer membrane of the target bacterial cell is solubilized, the MIC of the endolysin was not able to be determined by the microdilution checkerboard test. We therefore aimed to detect any decrease in the colistin MICs when used in combination with endolysin ElyA1. The addition of endolysin ElyA1 yielded a fourfold reduction in the colistin MICs in four of the six strains tested (*A. baumannii* GMA001 and PON001, *P. aeruginosa* AUS531 and *K. pneumoniae* KP17) (Figure 4). By contrast, only a twofold reduction in the colistin MIC was observed with *P. aeruginosa* AUS601 and no decrease with *K. pneumoniae* KP16. The latter was consistent with the lack of enzymatic activity observed in the antibacterial assays (Figure 4). Finally, no antimicrobial activity was detected when the combination was tested in the colistin resistant isolates (data not shown).

**Figure 4.**
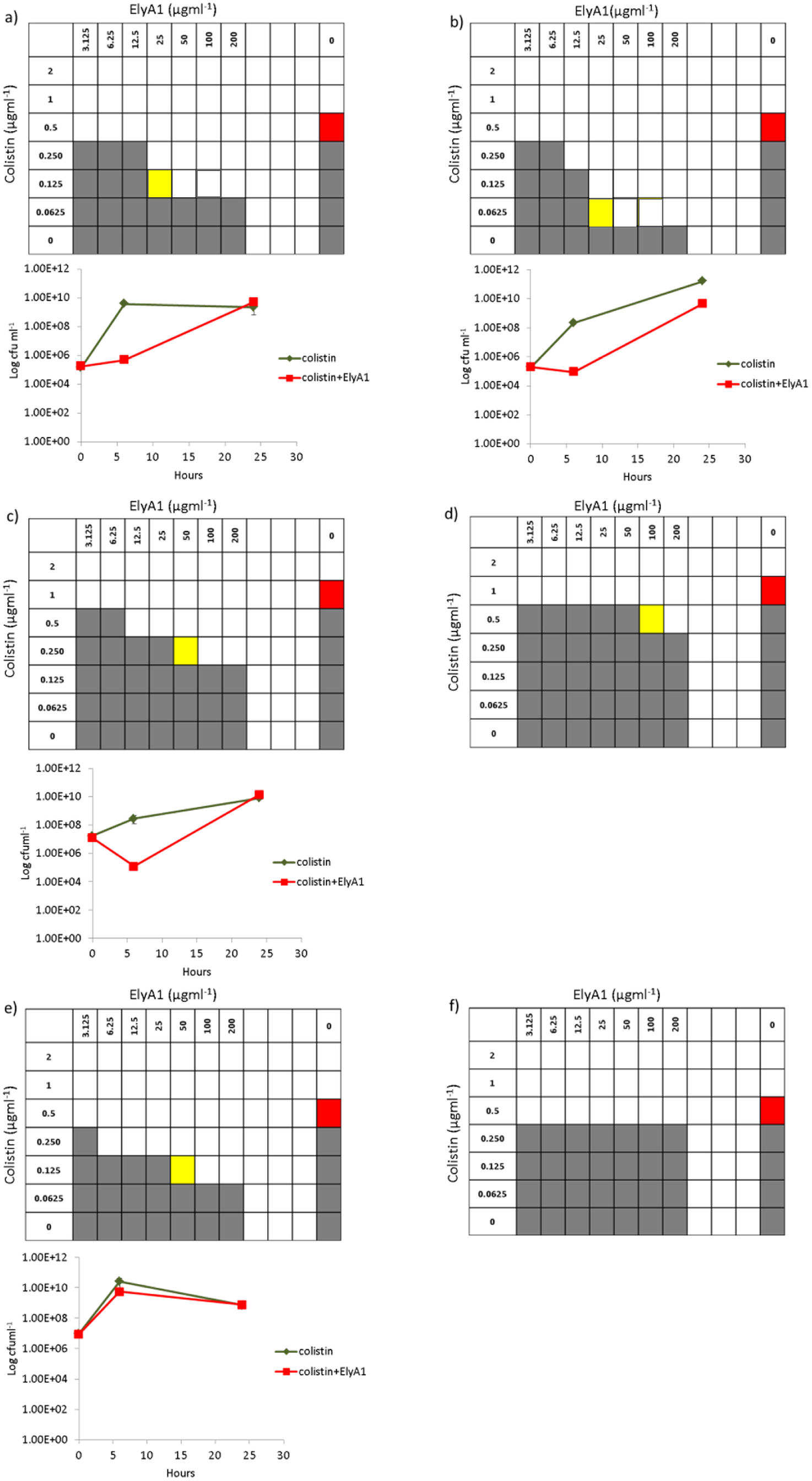
Bactericidal activity of colistin in combination with endolysin ElyA1 measured by MIC and time kill curves in *Acinetobacter baumannii* strains GMA001 (a) and PON001 (b); *Pseudomonas aeruginosa* strains AUS531(c) and AUS601(d); *Klebsiella pneumoniae* strains KP17(e) and KP16 (f). The time kill curves were only constructed for strains in which there was a fourfold reduction in colistin MICs (red square) when used in combination with the endolysin ElyA1(yellowsquare).

The results of the time kill curve assay confirmed the results obtained by the microdilution checkerboard test (Figure 4). A 2 log reduction was shown in the growth of both the *A. baumannii* strains and *P. aeruginosa* AUS531 after 6 hours in the culture with 1/4 of colistin and endolysin ElyA1, indicating a synergetic reaction between colistin and endolysin ElyA1. By contrast, no reduction was observed in the *K. pneumoniae* KP17 culture.

### Mortality in the *in vivo Galleria mellonella* Model

Larvae of the wax moth were infected with the clinical strain of *A. baumannii* PON001. When infected larvae were treated with colistin (¼MIC) in combination with endolysin ElyA1 (25 μg/ml), a significative increase (P<0.05) in the survival was observed in the treated larvae respect to those treated only with colistin (¼ MIC) (Figure 5a). When colistin (¼ MIC) was combined with ElyA2 (25 μg/ml) no significative differences (P>0.05) were obtained when compared with the treatment with colistin, as no muralytic activity was previously detected with ElyA2 (Figure 5b).

**Figure 5.**
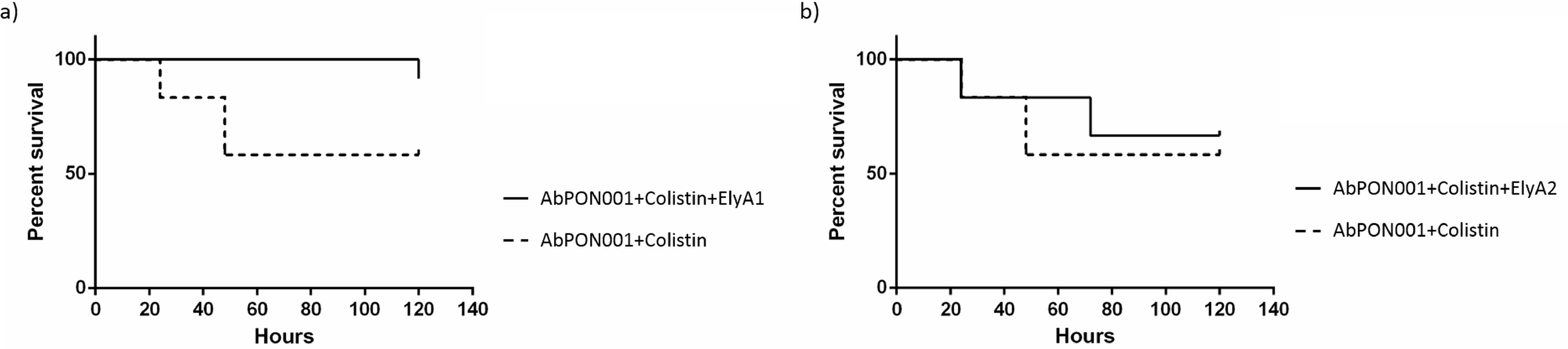
Survival curves of *G. mellonella* larvae infected with *A. baumannii* clinical strain PON001. a) Treated with colistin (1/4 MIC) and with colistin (1/4 MIC) combined with endolysin ElyA1 (25μg/ml); b) Treated with colistin (1/4 MIC) and with colistin (1/4 MIC) combined with endolysin ElyA2 (25μg/ml). This experiment was carried out with an adequate survival control.

## DISCUSSION

The discovery and development of novel antimicrobial agents to treat infections caused by the “priority” group of pathogens has occurred in the medical and research community (2).

Enzybiotics have become a focus of attention of many research groups worldwide. Endolysins, one type of enzybiotic, are species or genus-specific enzymes that act by hydrolysing the peptidoglycan layer of the bacterial cell wall. There are no reports of bacteria developing resistance to endolysins, which is a great problem in both antibiotic therapy and also in phage therapy (16). Moreover, endolysins have been recognized by the United States of America in the National Action Plan for Combating Antibiotic-resistant Bacteria (33), which identified the use of “phage-derived lysins to kill specific bacteria while preserving the microbiota” as a key strategy to reduce the development of antimicrobial resistance (34) due to the absence of toxicity in human cells (35).

The outer membrane of Gram-negative bacteria acts as a barrier preventing access of many endolysins to their natural target, the peptidoglycan layer. This problem has been approached by different strategies, including solubilization of the outer membrane with EDTA, modification of the endolysin PGs by deletion or substitution and by the development of fusion proteins such Artilysin-175, made by fusing the endolysin with a peptide to enable the enzyme to pass through the outer membrane which has been a successful strategy (18, 36, 37). In *P.aeruginosa* strains, Briers et al. (18) worked in the development of Art-175 constituted by antimicrobial peptide (AMP) sheep myeloid 29-amino acid peptide (SMAP-29) fused with endolysin KZ144. Art-175 showed muralytic activity in *P.aeruginosa* isolate and continuous exposure to Art-175 did not elicit the development of resistance. On its own, SMAP-29 is cytotoxic to mammalian cells (38); however, Art-175 exhibited little toxicity in L-292 mouse connective tissue.

As a new strategy we combined the membrane destabilizing effect of the colistin (“last-line” treatment for these bacteria due to the concerns about its nephrotoxicity and neurotoxicity (39)), which is a cationic peptide used as an active outer membrane agent, and two endolysins identified by our research group that belong to a lysozyme-like family (GH108-PG3) never before used as antimicrobial treatment.

In this work we identified two endolysins, ElyA1 and ElyA2, obtained from *A. baumannii* bacteriophage Ab1051Φ and Ab105Φ2, available in a collection of clinical strains of *A. baumannii* isolated during the II Spanish Multicentre Study GEIH/REIPI-*A.baumannii* 2000-2010 (Accession number, Genbank Umbrella Bioproject PRJNA422585) (26, 40).

Endolysins ElyA1 and ElyA2 are lysozyme-like proteins with a catalytic domain and a cell wall binding domain (CBD), responsible for recognition of the cell surface ligands and affinity for the bacterial substrate (41, 42). This structure is most commonly found in endolysins from bacteriophages that target Gram-positive bacteria. However, the PG_3 domain present in endolysins ElyA1 and ElyA2 has been identified in some Gram-negative bacteria and in a group of nine endolysins isolated from bacteriphages of *A. baumannii*; the domain shows high homology with ElyA1 and belongs to the same family (Figure 1) (20, 40, 43). Our molecular features and comparative genomes results in bacteriophage endolysins were confirmed through the work of Oliveira et al (44).

Furthermore, the bacteriophages from which these endolysins were isolated, Ab1051Φ and Ab105Φ2, occur in a large number of clinical isolates of *A. baummanii* (26). The cell wall binding domain has been shown to be responsible for the specificity and affinity of the endolysins for its substrate (43). However, we observed that endolysin ElyA1 displays a broader spectrum of activity against strains of *A. baumannii* and also many strains of *P. aeruginosa* belonging to the same order, *Pseudomonadales*, and to lesser extent some strains of *K. pneumoniae* from another gammaproteobacterial order, *Enterobacteriales.* In this case, the target of endolysin ElyA1, identified in Peptidoglycan binding domains (PGs) as a D-Asn (43), is probably conserved among the *Pseudomonadales*, thus explaining the broad spectrum of action of this enzyme. Interestingly, we were not able to detect muralytic activity in endolysin ElyA2 because this enzyme induces the aggregation of the cells *in vitro*, probably preventing the activity of the enzyme over all the cells.

In the present study, we used the cationic polymyxin antibiotic colistin to overcome the impenetrability of the outer membrane to endolysin ElyA1. Colistin disturbs the outer membrane via an electrostatic interaction with lipopolysaccharides and phospholipids present in the outer membrane (4). The synergistic effect of endolysin LysABP-01 (Lysozyme-like protein from the GH19 family) and colistin on *A. baumannii* has previously been described (25). Although endolysin ElyA1 does not display exogenous activity, because of its inability to cross the outer membrane, this problem was largely overcome when the enzyme was used in combination with colistin, and the antimicrobial activity of the combined therapy was higher than for both substances used alone in all the tested strains except for those of *K. pneumoniae*. A reduction in the colistin MIC of at least fourfold occurred for all of the *A. baumannii* strains tested and for *P. aeruginosa* strain AUS531, and a corresponding twofold reduction was observed for *P. aeruginosa* strain AUS601. A reduction in the colistin MIC was also obtained for the *K. pneumoniae* strain KP17, the most susceptible to endolysin ElyA1. The increased antimicrobial activity with endolysin ElyA1 and colistin was confirmed with the almost 3 log reduction in growth after 6 h in all strains tested, except KP17 *K. pneumoniae*. Growth of the culture reached the same level as in the control after 24 h, probably due to degradation of the enzyme and colistin, as also reported by Thummeepak *et al.* (22). In all of the strains tested the reduction in the colistin MIC was consistent with the muralytic activity of endolysin ElyA1 obtained for those strains. No antimicrobial activity was observed when this assay was conducted in colistin-resistant strains, probably because of the inability of the enzyme to access the peptidoglycan layer, as the necessary destabilization of the outer membrane by the colistin was not produced in these isolates. However, there are several resistance mechanisms to colistin described which include those in which the lipopolysaccharide is modified or is not produced preventing the joint of the colistin to the outer membrane, but there are other mechanisms in the literature which include efflux pumps described in *A. baumannii* and inhibition of respiratory enzymes as NADH oxidase in Gram positives as *Bacillus spp* and NADH quinone oxidoreductase in *E. coli.* It is probable that the activity of the enzyme would be higher in the bacteria with colistin resistance mechanisms different from that who modify the lipopolysaccharides (45–51). Future studies will be done over a range of different colistin resistant bacteria in order to reduce the MIC for colistin in combination with endolysin ElyA1.

The results obtained *in vitro* were confirmed with the assays done *in vivo*, as the survival of the infected larvae of *G. mellonella* was higher when the worms were treated with a combination of a reduced fourfold MIC of colistin and endolysin ElyA1 than with colistin alone. As a control the same assay was performed with endolysin ElyA2, in which the muralytic activity was not detected, and no differences were observed with the treatment with colistin.

In conclusion, this is the first study *in vitro* and *in vivo* (*G.mellonella*) where colistin is combined with endolysin ElyA1 (Glycosyde hydrolase 108 superfamily) against clinical MDR pathogens, which may enable the concentration of colistin used in antimicrobial treatments to be reduced, thus also reducing the toxic side effects of the antibiotic. The broad spectrum of action of endolysin ElyA1 would enable more MDR Gram-negative bacteria to be included as targets for this combined antimicrobial treatment.

## ACKNOWLEDGEMENTS

This study was funded by grant PI16/01163 awarded to M. Tomás within the State Plan for R+D+I 2013-2016 (National Plan for Scientific Research, Technological Development and Innovation 2008-2011) and co-financed by the ISCIII-Deputy General Directorate for Evaluation and Promotion of Research - European Regional Development Fund “A way of Making Europe” and Instituto de Salud Carlos III FEDER, Spanish Network for the Research in Infectious Diseases (REIPI, RD16/0016/0001, RD16/0016/0002, RD16/0016/0004, RD16/0016/0006, RD16/0016/0008, and RD16/0016/0011) and by the Study Group on Mechanisms of Action and Resistance to Antimicrobials, GEMARA (SEIMC, http://www.seimc.org/). M. Tomás was financially supported by the Miguel Servet Research Programme (SERGAS and ISCIII). R. Trastoy and L. Fernández-García were financially supported by respectively an SEIMC grant and predoctoral fellowship from the Xunta de Galicia (GAIN, Axencia de Innovación). TJK is supported by a National Health and Medical Research Council Early Career Fellowship (GNT1088448).

## TRANSPARENCY DECLARATIONS

All authors declare no conflict of interest

## REFERENCES

1. Bragg RR, Meyburgh CM, Lee JY, et al. Potential Treatment Options in a Post-antibiotic Era. Adv Exp Med Biol. 2018;1052:51–61. Bragg RR, Meyburgh CM, Lee JY, Coetzee M. 2018. Potential Treatment Options in a Post-antibiotic Era. Adv Exp Med Biol 1052:51–61.

2. Tacconelli E, Carrara E, Savoldi A, Harbarth S, Mendelson M, Monnet DL, Pulcini C, Kahlmeter G, Kluytmans J, Carmeli Y, Ouellette M, Outterson K, Patel J, Cavaleri M, Cox EM, Houchens CR, Grayson ML, Hansen P, Singh N, Theuretzbacher U, Magrini N, Group WPPLW. 2018. Discovery, research, and development of new antibiotics: the WHO priority list of antibiotic-resistant bacteria and tuberculosis. Lancet Infect Dis 18:318–327.

3. Gai Z, Samodelov SL, Kullak-Ublick GA, Visentin M. 2019. Molecular Mechanisms of Colistin-Induced Nephrotoxicity. Molecules 24.

4. Ordooei Javan A, Shokouhi S, Sahraei Z. 2015. A review on colistin nephrotoxicity. Eur J Clin Pharmacol 71:801–10.

5. Hojckova K, Stano M, Klucar L. 2013. phiBIOTICS: catalogue of therapeutic enzybiotics, relevant research studies and practical applications. BMC Microbiol 13:53.

6. Oliveira H, Vilas Boas D, Mesnage S, Kluskens LD, Lavigne R, Sillankorva S, Secundo F, Azeredo J. 2016. Structural and Enzymatic Characterization of ABgp46, a Novel Phage Endolysin with Broad Anti-Gram-Negative Bacterial Activity. Front Microbiol 7:208.

7. Elbreki M, Ross RP, Hill C, O’Mahony J, McAuliffe O, Coffey A. 2014. Bacteriophages and Their Derivatives as Biotherapeutic Agents in Disease Prevention and Treatment. Journal of Viruses 2014:20.

8. Schmelcher M, Powell AM, Becker SC, Camp MJ, Donovan DM. 2012. Chimeric phage lysins act synergistically with lysostaphin to kill mastitis-causing *Staphylococcus aureus* in murine mammary glands. Appl Environ Microbiol 78:2297–305.

9. Fenton M, Ross P, McAuliffe O, O’Mahony J, Coffey A. 2010. Recombinant bacteriophage lysins as antibacterials. Bioeng Bugs 1:9–16.

10. Gu J, Zuo J, Lei L, Zhao H, Sun C, Feng X, Du C, Li X, Yang Y, Han W. 2011. LysGH15 reduces the inflammation caused by lethal methicillin-resistant *Staphylococcus aureus* infection in mice. Bioeng Bugs 2:96–9.

11. Oechslin F, Daraspe J, Giddey M, Moreillon P, Resch G. 2013. In vitro characterization of PlySK1249, a novel phage lysin, and assessment of its antibacterial activity in a mouse model of *Streptococcus agalactiae* bacteremia. Antimicrob Agents Chemother 57:6276–83.

12. Loeffler JM, Djurkovic S, Fischetti VA. 2003. Phage Lytic Enzyme Cpl-1 as a Novel Antimicrobial for Pneumococcal Bacteremia. Infection and Immunity 71:6199–6204.

13. Lewinsohn DM, Leonard MK, LoBue PA, Cohn DL, Daley CL, Desmond E, Keane J, Lewinsohn DA, Loeffler AM, Mazurek GH, O’Brien RJ, Pai M, Richeldi L, Salfinger M, Shinnick TM, Sterling TR, Warshauer DM, Woods GL. 2017. Official American Thoracic Society/Infectious Diseases Society of America/Centers for Disease Control and Prevention Clinical Practice Guidelines: Diagnosis of Tuberculosis in Adults and Children. Clin Infect Dis 64:111–115.

14. Lood R, Raz A, Molina H, Euler CW, Fischetti VA. 2014. A highly active and negatively charged *Streptococcus pyogenes* lysin with a rare D-alanyl-L-alanine endopeptidase activity protects mice against streptococcal bacteremia. Antimicrob Agents Chemother 58:3073–84.

15. Gilmer DB, Schmitz JE, Euler CW, Fischetti VA. 2013. Novel bacteriophage lysin with broad lytic activity protects against mixed infection by Streptococcus pyogenes and methicillin-resistant *Staphylococcus aureus*. Antimicrob Agents Chemother 57:2743–50.

16. Roach DR, Donovan DM. 2015. Antimicrobial bacteriophage-derived proteins and therapeutic applications. Bacteriophage 5:e1062590.

17. Lai M-J, Lin N-T, Hu A, Soo P-C, Chen L-K, Chen L-H, Chang K-C. 2011. Antibacterial activity of *Acinetobacter baumannii* phage ϕAB2 endolysin (LysAB2) against both Gram-positive and Gram-negative bacteria. Applied Microbiology and Biotechnology 90:529–539.

18. Briers Y, Walmagh M, Grymonprez B, Biebl M, Pirnay JP, Defraine V, Michiels J, Cenens W, Aertsen A, Miller S, Lavigne R. 2014. Art-175 is a highly efficient antibacterial against multidrug-resistant strains and persisters of Pseudomonas aeruginosa. Antimicrob Agents Chemother 58:3774–84.

19. Briers Y, Walmagh M, Van Puyenbroeck V, Cornelissen A, Cenens W, Aertsen A, Oliveira H, Azeredo J, Verween G, Pirnay JP, Miller S, Volckaert G, Lavigne R. 2014. Engineered endolysin-based “Artilysins” to combat multidrug-resistant gram-negative pathogens. MBio 5:e01379–14.

20. Lood R, Winer BY, Pelzek AJ, Diez-Martinez R, Thandar M, Euler CW, Schuch R, Fischetti VA. 2015. Novel phage lysin capable of killing the multidrug-resistant gram-negative bacterium *Acinetobacter baumannii* in a mouse bacteremia model. Antimicrob Agents Chemother 59:1983–91.

21. Briers Y, Walmagh M, Lavigne R. 2011. Use of bacteriophage endolysin EL188 and outer membrane permeabilizers against *Pseudomonas aeruginosa*. J Appl Microbiol 110:778–85.

22. Thummeepak R, Kitti T, Kunthalert D, Sitthisak S. 2016. Enhanced Antibacterial Activity of *Acinetobacter baumannii* Bacteriophage ØABP-01 Endolysin (LysABP-01) in Combination with Colistin. Front Microbiol 7:1402.

23. Ibarra-Sánchez LA, Van Tassell ML, Miller MJ. 2018. Antimicrobial behavior of phage endolysin PlyP100 and its synergy with nisin to control *Listeria monocytogenes* in Queso Fresco. Food Microbiol 72:128–134.

24. Letrado P, Corsini B, Díez-Martínez R, Bustamante N, Yuste JE, García P. 2018. Bactericidal synergism between antibiotics and phage endolysin Cpl-711 to kill multidrug-resistant pneumococcus. Future Microbiol 13:1215–1223.

25. Kim NH, Park WB, Cho JE, Choi YJ, Choi SJ, Jun SY, Kang CK, Song KH, Choe PG, Bang JH, Kim ES, Park SW, Kim NJ, Oh MD, Kim HB. 2018. Effects of Phage Endolysin SAL200 Combined with Antibiotics on *Staphylococcus aureus* Infection. Antimicrob Agents Chemother 62.

26. López M, Rueda A, Florido JP, Blasco L, Fernández-García L, Trastoy R, Fernández-Cuenca F, Martínez-Martínez L, Vila J, Pascual A, Bou G, Tomas M. 2018. Evolution of the Quorum network and the mobilome (plasmids and bacteriophages) in clinical strains of *Acinetobacter baumannii* during a decade. Sci Rep 8:2523.

27. López-Causapé C, Sommer LM, Cabot G, Rubio R, Ocampo-Sosa AA, Johansen HK, Figuerola J, Cantón R, Kidd TJ, Molin S, Oliver A. 2017. Evolution of the *Pseudomonas aeruginosa* mutational resistome in an international Cystic Fibrosis clone. Sci Rep 7:5555.

28. Esteban-Cantos A, Aracil B, Bautista V, Ortega A, Lara N, Saez D, Fernández-Romero S, Pérez-Vázquez M, Navarro F, Grundmann H, Campos J, Oteo J, Network E-S. 2017. The Carbapenemase-Producing *Klebsiella pneumoniae* Population Is Distinct and More Clonal than the Carbapenem-Susceptible Population. Antimicrob Agents Chemother 61.

29. Schmitz JE, Schuch R, Fischetti VA. 2010. Identifying active phage lysins through functional viral metagenomics. Appl Environ Microbiol 76:7181–7.

30. Sopirala MM, Mangino JE, Gebreyes WA, Biller B, Bannerman T, Balada-Llasat JM, Pancholi P. 2010. Synergy testing by Etest, microdilution checkerboard, and time-kill methods for pan-drug-resistant *Acinetobacter baumannii*. Antimicrob Agents Chemother 54:4678–83.

31. Peleg AY, Jara S, Monga D, Eliopoulos GM, Moellering RC, Mylonakis E. 2009. Galleria mellonella as a model system to study *Acinetobacter baumannii* pathogenesis and therapeutics. Antimicrob Agents Chemother 53:2605–9.

32. Yang H, Chen G, Hu L, Liu Y, Cheng J, Li H, Ye Y, Li J. 2015. In vivo activity of daptomycin/colistin combination therapy in a Galleria mellonella model of *Acinetobacter baumannii* infection. Int J Antimicrob Agents 45:188–91.

33. U.S. Office of the Press Secretary: Washington D, USA,. 2015. The White House. National Action Plan for Combating Antibiotic-Resistant Bacteria. Interagency Task Force for Combating Antibiotic-Resistant Bacteria;.

34. Love MJ, Bhandari D, Dobson RCJ, Billington C. 2018. Potential for Bacteriophage Endolysins to Supplement or Replace Antibiotics in Food Production and Clinical Care. Antibiotics (Basel) 7.

35. Nguyen S, Baker K, Padman BS, Patwa R, Dunstan RA, Weston TA, Schlosser K, Bailey B, Lithgow T, Lazarou M, Luque A, Rohwer F, Blumberg RS, Barr JJ. 2017. Bacteriophage Transcytosis Provides a Mechanism To Cross Epithelial Cell Layers. MBio 8.

36. Mao J, Schmelcher M, Harty WJ, Foster-Frey J, Donovan DM. 2013. Chimeric Ply187 endolysin kills *Staphylococcus aureus* more effectively than the parental enzyme. FEMS Microbiol Lett 342:30–6.

37. Mayer MJ, Garefalaki V, Spoerl R, Narbad A, Meijers R. 2011. Structure-based modification of a *Clostridium difficile*-targeting endolysin affects activity and host range. J Bacteriol 193:5477–86.

38. Kosikowska P, Lesner A. 2016. Antimicrobial peptides (AMPs) as drug candidates: a patent review (2003-2015). Expert Opin Ther Pat 26:689–702.

39. Da Silva GJ, Domingues S. 2017. Interplay between Colistin Resistance, Virulence and Fitness in *Acinetobacter baumannii*. Antibiotics (Basel) 6.

40. López M, Rueda A, Florido JP, Blasco L, Gato E, Fernández-García L, Martínez-Martínez L, Fernández-Cuenca F, Pachón J, Cisneros JM, Garnacho-Montero J, Vila J, Rodríguez-Baño J, Pascual A, Bou G, Tomás M. 2016. Genomic Evolution of Two *Acinetobacter baumannii* Clinical Strains from ST-2 Clones Isolated in 2000 and 2010 (ST-2_clon_2000 and ST-2_clon_2010). Genome Announc 4.

41. Veiga-Crespo P, Ageitos JM, Poza M, Villa TG. 2007. Enzybiotics: a look to the future, recalling the past. J Pharm Sci 96:1917–24.

42. Yang H, Yu J, Wei H. 2014. Engineered bacteriophage lysins as novel anti-infectives. Front Microbiol 5:542.

43. Jarábková V, Tišáková L, Godány A. 2015. Phage Endolysin: A Way To Understand A Binding Function Of C-Terminal Domains A Mini Review. 14:117.

44. Oliveira H, Melo LD, Santos SB, Nóbrega FL, Ferreira EC, Cerca N, Azeredo J, Kluskens LD. 2013. Molecular aspects and comparative genomics of bacteriophage endolysins. J Virol 87:4558–70.

45. Deris ZZ, Akter J, Sivanesan S, Roberts KD, Thompson PE, Nation RL, Li J, Velkov T. 2014. A secondary mode of action of polymyxins against Gram-negative bacteria involves the inhibition of NADH-quinone oxidoreductase activity. J Antibiot (Tokyo) 67:147–51.

46. Cheah SE, Johnson MD, Zhu Y, Tsuji BT, Forrest A, Bulitta JB, Boyce JD, Nation RL, Li J. 2016. Polymyxin Resistance in *Acinetobacter baumannii:* Genetic Mutations and Transcriptomic Changes in Response to Clinically Relevant Dosage Regimens. Sci Rep 6:26233.

47. Ni W, Li Y, Guan J, Zhao J, Cui J, Wang R, Liu Y. 2016. Effects of Efflux Pump Inhibitors on Colistin Resistance in Multidrug-Resistant Gram-Negative Bacteria. Antimicrob Agents Chemother 60:3215–8.

48. Lin MF, Lin YY, Lan CY. 2017. Contribution of EmrAB efflux pumps to colistin resistance in *Acinetobacter baumannii*. J Microbiol 55:130–136.

49. Verma P, Maurya P, Tiwari M, Tiwari V. 2017. In-silico interaction studies suggest RND efflux pump mediates polymyxin resistance in *Acinetobacter baumannii*. J Biomol Struct Dyn:1–9.

50. Poirel L, Jayol A, Nordmann P. 2017. Polymyxins: Antibacterial Activity, Susceptibility Testing, and Resistance Mechanisms Encoded by Plasmids or Chromosomes. Clin Microbiol Rev 30:557–596.

51. Machado D, Antunes J, Simões A, Perdigão J, Couto I, McCusker M, Martins M, Portugal I, Pacheco T, Batista J, Toscano C, Viveiros M. 2018. Contribution of efflux to colistin heteroresistance in a multidrug resistant *Acinetobacter baumannii* clinical isolate. J Med Microbiol 67:740–749.

